# Planar cell polarity: the *prickle* gene acts independently on both the Ds/Ft and the Stan Systems

**DOI:** 10.1101/212167

**Authors:** José Casal, Beatriz Ibáñez-Jiménez, Peter A. Lawrence

## Abstract

Epithelial cells are polarised within the plane of the epithelium, forming oriented structures whose coordinated and consistent polarity (planar cell polarity, PCP) relates to the principal axes of the body or organ. In *Drosophila* at least two separate molecular systems generate and interpret intercellular polarity signals: Dachsous/Fat, and the “core” or Stan system. Here we study the *prickle* gene and its protein products Prickle and Spiny leg. Much research on PCP has focused on the asymmetric localisation of core proteins in the cell and as a result *prickle* was placed in the heart of the Stan system. Here we ask if this view is correct and how the *prickle* gene relates to the two systems. We find that *prickle* can affect, separately, both systems — however, neither Pk nor Sple are essential components of the Ds/Ft or the Stan system, nor do they act as a functional link between the two systems.

## Introduction

Planar cell polarity (PCP) refers to a property that all, or most, epithelial cells have — they are coordinately oriented in the plane of the epithelial sheet and, sometimes, they demonstrate this by forming oriented structures. These oriented structures can be cell organelles such as cilia, or multicellular organs such as mammalian hairs (Tree et al., 2002a; Wang and Nathans, 2007; Goodrich and Strutt, 2011; Butler and Wallingford, 2017). *Drosophila* has been used to identify most of the genes involved in PCP and has proved the most amenable of all animals for elucidating its mechanisms. Most studies have asked where PCP gene products are localised in the cell and asked how these localisations relate to the propagation of polarity from cell to cell. Here we use genetic methods to investigate the function of the *prickle* (*pk*), a PCP gene.

The *pk* gene produces two homologous transcripts that encode the Pk and Sple proteins; both proteins contain protein-protein binding LIM domains, they differ in the N terminus (Gubb et al., 1999) and have sequence elements conserved to vertebrates. In vertebrates, syndromes due to *pk* mutations have been identified but it is not clear if they reveal genuine PCP functions of the *pk* genes (Tissir and Goffinet, 2013). In flies, in the absence of the *pk* gene, the polarities of bristles and hairs are altered and/or disordered over large areas of the cuticle. In the abdomen, when the *pk* isoform is overexpressed everywhere, polarity of the anterior (A) compartment is almost entirely reversed while the posterior (P) compartment is normal. By contrast, when the *sple* isoform is overexpressed everywhere, polarity of the P compartment is completely reversed while the A compartment is normal. These findings led to the hypothesis that the *pk* gene functions in the wildtype to turn around or “rectify” polarity in order to coordinate PCP over regions of the fly (Lawrence et al., 2004). These results are consistent with the hypothesis that Pk and Sple have similar functions in PCP, the local outcome depending on the distribution of both proteins and varied regional responses to them (Gubb and Garcia-Bellido, 1982). For example, in the wildtype, of the two proteins, Pk is found to be the most effective agent in the wing, in the P compartment of the abdomen and the posterior part of the thorax while Sple is thought to predominate in the A compartment of the abdomen and anterior region of the thorax (Gubb et al., 1999; Ayukawa et al., 2014; Merkel et al., 2014; Ambegaonkar and Irvine, 2015).

One set of proteins constitute the “core system” of PCP. Because of the central importance of Starry night (Stan) to function of this system we renamed it the “Stan system” (Lawrence et al., 2004). Proteins classified as members of the Stan system include Stan, Fz, Vang Gogh (Vang), Dishevelled (Dsh), Diego (Dgo) and Pk/Sple (for a review see Goodrich and Strutt, 2011; Adler, 2012; Butler and Wallingford, 2017). Some of these proteins are asymmetrically distributed: in the wing, Pk is enriched on or near the proximal membrane of each cell (Tree et al., 2002b), while Frizzled (Fz) is localised distally (Strutt, 2001). The localisation of these proteins is mutually dependent; when one protein is removed, the others become evenly distributed around the cell periphery (reviewed in Strutt and Strutt, 2009). The current view is that Dsh, Pk and Fz interact with each other to amplify their asymmetric localisation within the cell and thereby consolidate its polarity. Groups of proteins associate in separate complexes on different sides of the wing cell and are associated with each other across the cell membrane, proximally or distally, as a response to an upstream signal (reviewed in Klein and Mlodzik, 2005).

Some argue that PCP is produced by three tiers of gene activity (Tree et al., 2002a; Tree et al., 2002b; Yang et al., 2002; Klein and Mlodzik, 2005; Strutt and Strutt, 2005; Strutt and Strutt, 2007; Axelrod, 2009) in which asymmetrical distribution of the protocadherins Dachsous (Ds) and Fat (Ft) provides an initial cue to orient the Stan system which amplifies the signal to polarise effector functions. Recently it has been posited that the *pk* gene intervenes between the polarising information specified by the direction of the gradients of Ds/Ft activity and its interpretation by the Stan system. These articles (Hogan et al., 2011; Ayukawa et al., 2014; Olofsson et al., 2014; Ambegaonkar and Irvine, 2015) also support earlier conclusions that Pk and Sple act discordantly on polarity output in different tissues and have improved the evidence that changes in the levels of Pk and/or Sple can turn around the orientation of polarised structures (Ayukawa et al., 2014). Moreover, they present arguments that Sple is the main component of a molecular link between the Ds/Ft and Stan systems.

But, some earlier findings are inconsistent with the hypothesis that Pk/Sple acts as a link:

1. The two systems can work independently of each other, at least in the abdomen (Casal et al., 2006), meaning that neither the Ds/Ft system nor the Stan system need each other to polarise cells. If so, there would be no need for a general link between the two systems.
2. Genetic experiments argue that, functionally, Pk and Sple are not required for polarity signalling from cell to cell, a central aspect of PCP. Adler et al. (2000) found that a weak allele of *pk* did not block or inhibit but actually increased those local changes in cell polarity that are induced by clones mutant for other Stan system genes. Adler’s finding was confirmed when Lawrence et al. (2004) showed that complete loss of Pk and Sple increases polarisation by the Stan system genes; they proposed that the key molecules in the Stan system do not include Pk but are Stan, Fz and Vang. This conclusion was later supported by Strutt and Strutt (2007) who presented further evidence that Dsh, Pk and Dgo are not needed for the propagation of polarity from cell to cell.

Here we undertake genetic experiments aimed at clarifying the function of Pk/Sple in the wildtype abdomen, particularly in relation to the two systems. We conclude that Pk and Sple are not essential components of either system nor are they essential components of a link between the two systems. We add to evidence that the Ds/Ft system acts independently of the Stan system and, surprisingly, provide data that Pk and Sple can rectify the output of the Ds/Ft system, even in the absence of a functioning Stan system. Pk and Sple also, separately, affect the output of the Stan system; they alter how far the polarising signal can spread.

## Results

### Explaining terms and methods

We make genetically marked clones of cells of different genotypes to ask how the two different genetic systems, the Ds/Ft system and the Stan system, define cell polarity in the anterior (A) and posterior compartments (P) of the adult abdomen. We assay function of the Ds/Ft system by the ability of “sending cells” in clones that, say, overexpress *ft*, to change the polarity of “receiving cells”. Such responses that are induced in cells nearby the clone are defined as “non-autonomous”. As a result, hairs and bristles around the clones may point “inwards” or “outwards”, that is, in or away from the clone. For the Ds/Ft system, Ds, Ft and Dachs (D) are each essential; however removal of only Ds or Ft causes misdistribution of the D protein in each cell (Ambegaonkar et al., 2012; Pan et al., 2013), leading to an adventitious phenotype of whorly polarity (Ambegaonkar et al., 2012; Lawrence and Casal, 2013). Therefore the cleanest way to break the Ds/Ft system completely and persuasively is to remove D as well as Ds or Ft. To break the Stan system we remove Stan; *stan^-^* cells cannot send or receive signals, for example *stan^-^* receiving cells cannot respond to cells that overexpress *fz* (Genotype 1) even when those sending cells also express *stan^-^* (Lawrence et al., 2004; Casal et al., 2006). Using these functional assays we ask whether and how Pk and Sple cooperate with the Ds/Ft and the Stan systems.

Figure 1 acts as a summary of, and a guide to, all the experiments and results.

**Figure 1.**
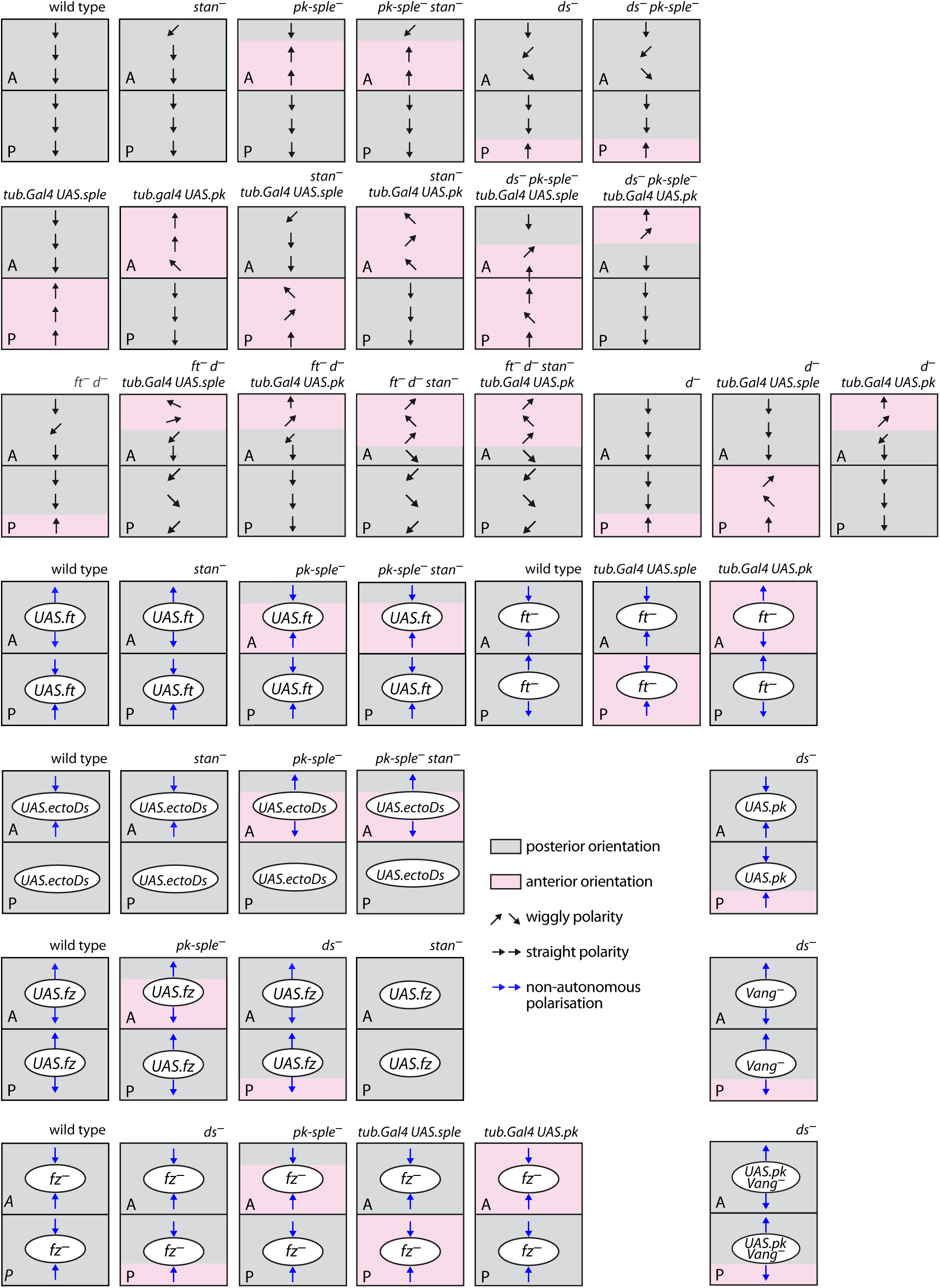
A guide to all the experiments. A summary of the results showing the polarities of hairs in the two abdominal compartments plus the effects of clones on polarity. *UAS* indicates overexpression of the said gene in the clones, *tub.Gal4 UAS.x* indicates generalised expression of x. Grey areas show hairs pointing generally posteriorly; pink areas indicate mainly reversed polarity as is also indicated by the arrows.

### Do the Ds/Ft and Stan systems act independently in both A and P compartments?

It has been proposed that upstream polarity information specified by the Ds/Ft system is interpreted by the Stan system (Yang et al., 2002; Ma et al., 2003; Goodrich and Strutt, 2011; reviewed in Butler and Wallingford, 2017). Experiments in the adult abdomen showed that the non-autonomous effects on neighbouring cells by clones, for example, lacking *ft*, depended on the compartment. *ft^-^* clones in the A compartment made the surrounding cells point inwards towards the clone, while the same clones in the P compartment caused the surrounding cells to point outwards. We therefore argued that the gradients of Ds and Fj activities might have opposite slopes in the anterior (A) and the posterior (P) compartments (Casal et al., 2002). But if that were true then, because the hairs in both compartments point the same way in the wildtype, hair polarity cannot be a direct readout of the gradient slope of the Ds/Ft system. Experimental evidence provided a solution to this conundrum: perhaps Pk or Sple rectify the reading of a gradient in either the A or the P compartment so that all hairs point in the same direction (Lawrence et al., 2004). But, later experiments argued that the Stan system and the Ds/Ft system can act independently of each other (Casal et al., 2006; Lawrence et al., 2007; and more recently Brittle et al., 2012) —implying that rectification due to Pk and/or Sple does not alter a direct input from the Ds/Ft system into the Stan system but avoids dissonance between their independent inputs into PCP.

#### (i) clones affecting the Ds/Ft system function when the Stan system is broken

Previously we showed that clones affecting the Ds/Ft system could polarise cells non-autonomously and do so very well in the absence of a functioning Stan system. These studies were limited to the A compartments. Here we show that, in both the A and the P compartments, clones overexpressing *ft* polarise both wildtype cells (Genotype 2) and cells in which the Stan system is broken —we have used flies lacking *stan*, (Genotype 3) or, in the case of A clones, both *stan* and *fz* (Casal et al., 2006). In both cases the receiving cells tend to point hairs outwards from the clone in the A compartments (Casal et al., 2006) and inwards in the P compartments (Figure 2 and Figure 3). Consistent with these results, clones overexpressing the extracellular domain of Ds also polarise both wildtype cells (Genotype 4) and cells in which the Stan system is broken (*stan^-^* Genotype 5) inwards in the anterior portion of A compartments (Casal et al., 2002; Casal et al., 2006). These clones are ineffective in the posterior parts of A compartments and in P compartments (Figure S3), probably because the activity of Ds is normally high in these regions (Casal et al., 2002).

**Figure 2.**
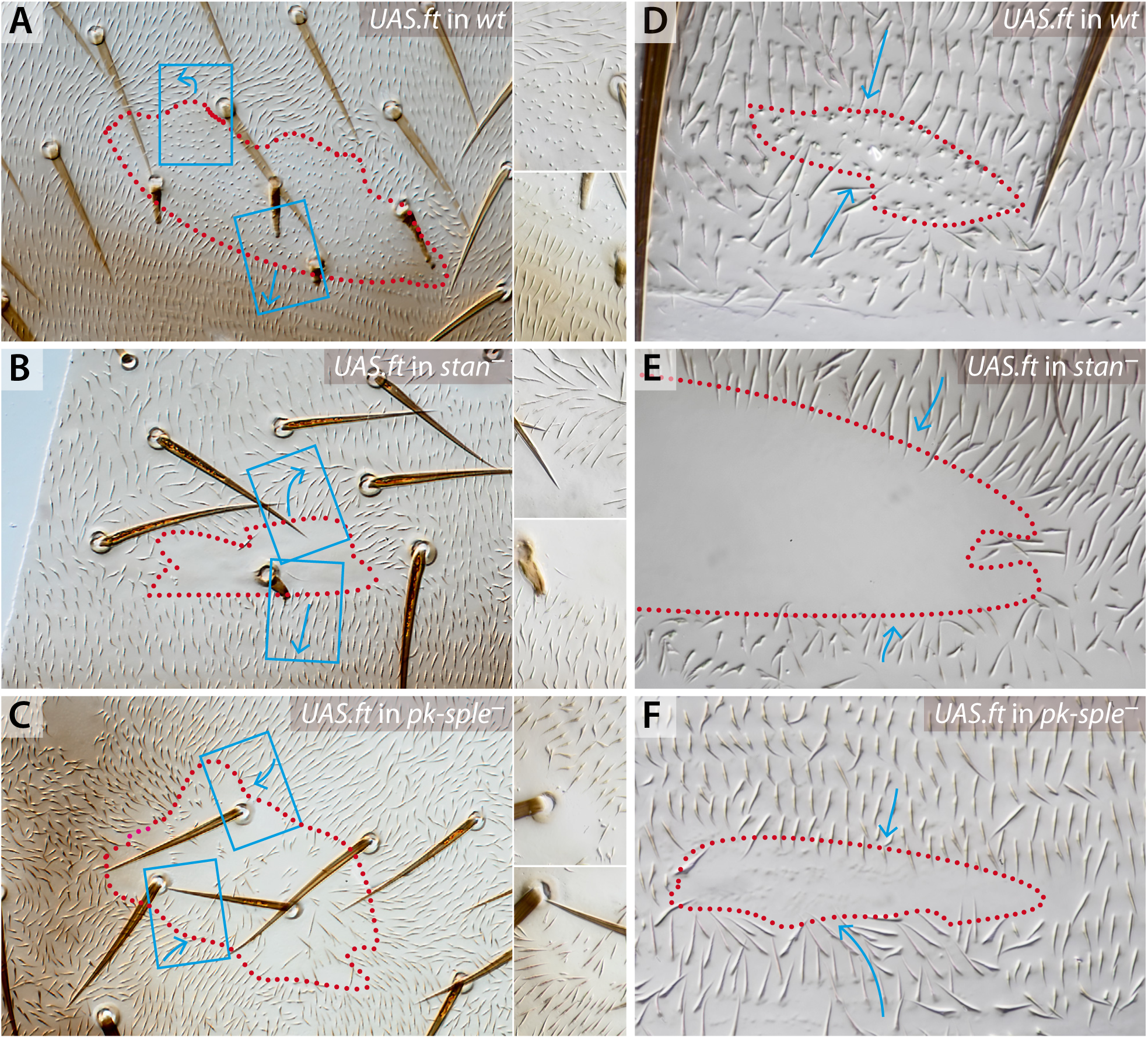
Clones that overexpress *ft* in various backgrounds. The receiving cells point outwards in the A compartments (A-B), inwards in P compartments (D-E) of *stan^-^* and wildtype cells. The response of *pk-sple^-^* cells is inwards in both the A and P compartments (C,F). For all figures, clones are outlined in red dots, blue boxes delimit the areas detailed at higher magnification, blue arrows indicate orientation of hairs. For images of clones expressing *fz* in the same backgrounds, see Figure S2.

**Figure 3.**
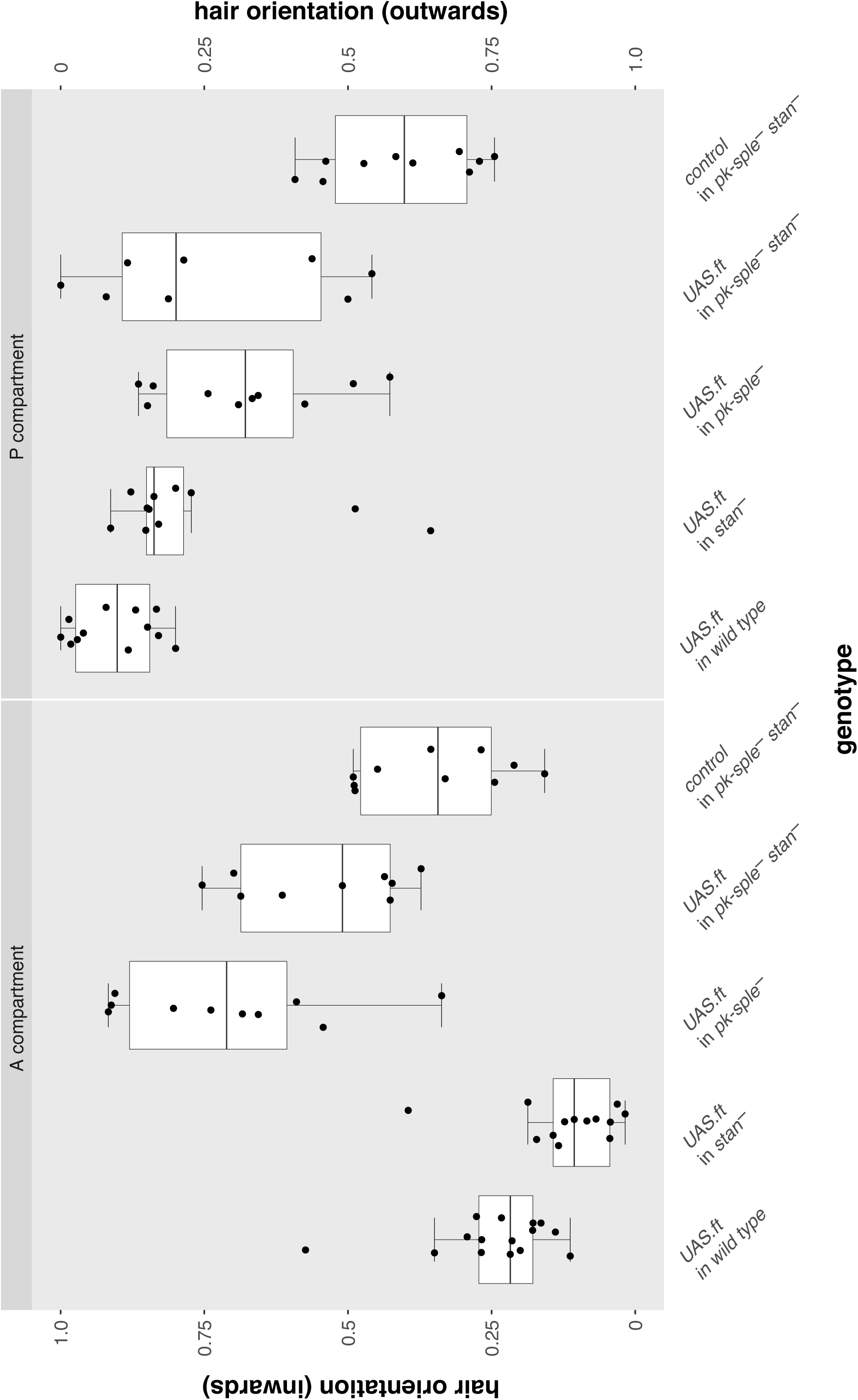
Effects of the *ft*-overexpressing clones in A and P compartments (cf Figure 2). The orientations of hairs immediately adjacent to each clone are counted and displayed in box plots, each dot represents the data from one clone. The responses range from all pointing inwards (top of the graph) to all pointing outwards (bottom). Breaking the Stan system (*stan^-^*) did not much affect any outcome, see Figure S3 for statistical analysis, confirming that the Ft/Ds system does not act through the Stan system. However removing *pk* and *sple* changed the sign of response in the A compartment. (Control clones Genotype 27).

#### (ii) clones affecting the Stan system function when the Ds/Ft system is broken

We can also look at the independence of the two systems by interfering with the Stan system in clones in a background in which the Ds/Ft system is broken. Clones that overexpress *fz*, in either the A or P compartments, normally turn the polarity of receiving cells to point outwards from the clone in A (Casal et al., 2006) and also in P (Genotype 6, Figure S2). They do the same in *ds^-^* flies but with a longer range (Genotype 7; Adler et al., 1998; Ma et al., 2003; Casal et al., 2006; Figure S1).

These and previous experiments have established that the two systems can act independently in both compartments of the abdomen; we now ask are Pk and Sple essential components of either the Ds/Ft or the Stan systems?

### How do Pk and Sple interact with each of the two systems?

#### (i) evidence from epistasis

*ds^-^* and *pk-sple^-^* flies differ in phenotype in the dorsal abdomen: the most useful difference is seen in the P compartment, where, in *ds^-^* flies, hairs in the anterior region of the P compartment are in whorls (probably due to the misdistribution of Dachs) but in the posterior region of the P compartment the hairs point directly anteriorward. By contrast, in *pk-sple^-^* flies, the entire P compartment has normal polarity (Lawrence et al., 2004). If Ds/Ft were to provide an upstream cue that is interpreted by the Stan system via Pk/Sple one would expect the double mutant *ds^-^ pk-sple^-^* to have a phenotype more similar to the single *pk-sple^-^* mutant than to the *ds^-^* mutant phenotype. However we find that, in the abdomen, *ds^-^ pk-sple^-^* flies (Genotype 8) are little different from *ds^-^* flies (Figure 1). It follows that the *ds* mutation is epistatic to a mutation that removes both *pk* and *sple* functions. Turning to interaction between the Stan system and Pk/Sple. When *pk-sple^-^ stan^-^* flies are compared to each single mutant, they differ from both, having an additive phenotype (Genotype 9, Genotype 10, and Genotype 11, Figure 1), implying that the two genes act independently. Taken together, these findings suggest that the *pk* gene acts entirely through the Ds/Ft system. However other results argue that Pk and Sple can also act independently of the Ds/Ft system (see below), and suggest instead that Pk and Sple act separately but differently on each of the two systems.

#### (ii) The Stan system functions well, both in cells that lack *pk* and *sple* and in cells that have *pk* or *sple* overexpressed

It has been proposed that Pk/Sple act as a link between Ds/Ft and the Stan system, rectifying the polarity output to ensure that the A and the P compartments have the same polarity. Here we test this link by making clones that alter the Stan system in *pk-sple^-^* mutant flies or in flies with ubiquitous overexpression of *pk* or *sple*.

1. In *pk-sple^-^* flies. In the abdomen of *pk-sple^-^* flies (Genotype 11), polarity of most of the A compartment is reversed, but the P compartment is normal. Clones of cells that overexpress *fz* (Genotype 12 or, alternatively, lack *fz*, Genotype 13) in such *pk-sple^-^* flies strongly polarise receiving cells in both A and P compartments; in both compartments the clones affect mutant receiving cells with the same sign as in wildtype receiving cells, that is outwards from the clones that overexpress *fz* and inwards towards clones that lack *fz*, independently of the prevailing polarity of the receiving cells (Figure S2). Thus, the Stan system does not need Pk or Sple to send polarity signals or to repolarise receiving cells (cf Lawrence et al., 2004).

2. When *pk* or *sple* are overexpressed. In flies in which either *sple* or *pk* are overexpressed, large areas of each abdominal segment show abnormal polarity. Nevertheless *fz^-^* clones made in these flies polarise receiving cells of both compartments inwards —as they do in wildtype flies—, independently of the prevailing polarity of those receiving cells (Genotype 14, Genotype 15, Figure 6). All these results are mutually consistent: they show that polarity changes induced by the Stan system do not require products of the *pk* gene, showing that Pk and Sple cannot be essential components of the Stan system in the wildtype.

3. However, the *fz^-^* clones do not behave exactly as they would in a wildtype background: absence or excess of Pk and Sple change the amount of polarisation caused by clones with altered amounts of Fz. In A compartments of the abdomen, clones of cells that lack *fz* alter polarity of surrounding wildtype cells. The number of rows of receiving cells affected, the range, varies with the amount of Pk and/or Sple protein: in *pk-sple^-^* flies (Genotype 13) the range of polarisation around the *fz^-^* clones or clones that contain excess *fz* (Figure 4 in Lawrence et al., 2004) is increased, resembling the increase in range observed when *fz* clones are induced in *ds^-^* flies (Genotype 16). Raising the level of the Pk isoform ubiquitously does not alter the range of polarisation surrounding *fz^-^* clones(Genotype 15), while when Sple levels are raised (Genotype 14), this range of polarisation is reduced (Figure S5). In the P compartments, we detected no effects on the range of repolarisation surrounding *fz^-^* clones; either in *pk-sple^-^* flies or when the levels of either Pk or Sple were increased (Figure S2 and Figure S5). These results add to evidence that the Stan system can function independently of Pk and Sple.

**Figure 4.**
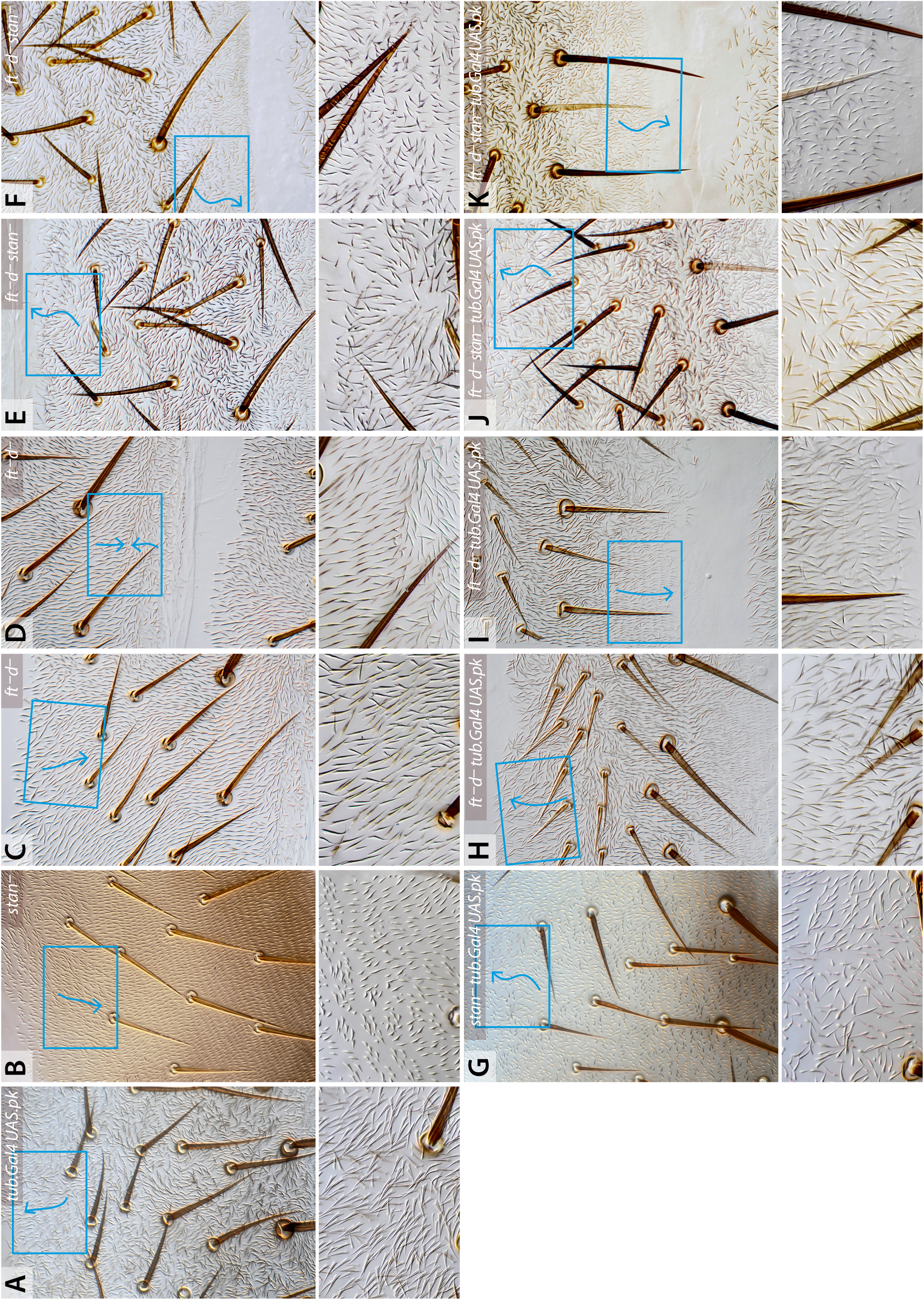
Effects of overexpressing *pk* on polarity of cells in which either the Stan system (*stan^-^*) or the Ds/Ft system is broken (ft^-^ d), or both are broken (*ft*^-^ *d^-^ stan^-^*). Background phenotypes for the A compartments (A, B, C and E) and for the P compartments (D, F). In the A compartments, generalised overexpression of *pk* changes the polarity of the anterior region of wildtype, *stan^-^* (Genotype 32) and *ft^-^ d^-^* cells (A, E and F). In the P compartments, the region that normally points anteriorly in *ft^-^ d^-^* points posteriorly (as in the wildtype) when *pk* is overexpressed (I). Ubiquitous Pk appears to have no effect on *ft^-^ d^-^ stan^-^* in either A or P compartments (J and K). Compare Figure S4 for expression of *pk* in *d^-^* flies.

#### (iii) The Ds/Ft system functions well, both in cells that lack Pk and Sple and in cells that have *pk* or *sple* overexpressed

If Pk/Sple act as an essential link between Ds/Ft and the Stan system one may expect that clones with an altered Ds/Ft system not to have any effect on hair polarity in clones lacking both Pk and Sple. However the results were striking and did not support the link hypothesis.

1. In *pk-sple^-^* flies. Clones of cells overexpressing *ft* repolarise receiving cells strongly, even if they lack Pk and Sple (Genotype 17). However it surprised us that in the largely reversed A compartment of the *pk-sple^-^* abdomen, the hairs around the clones point inwards (the opposite sign induced by such clones in the wildtype) and also inwards in the P compartment (the same sign as in wildtype, Figure 2). Clones overexpressing *ds* in *pk-sple^-^* flies (Genotype 18) act comparably, the hairs around such clones point outwards in A (the opposite sign induced by such clones in the wildtype) and outwards, but weakly, in the P compartment (the same sign as in wildtype, see Figure S3). Thus in clones of both genotypes, in the A compartments, the sign of the effect is the opposite from when such clones are made in the wildtype (Genotype 2 and Genotype 4). Nevertheless, in both these genotypes, in the P compartments, the sign of the polarising effect is the same as wildtype. Quantitation of overexpressing *ft* and *ds* clones confirms these results and also shows that these clones (in the A compartment) affect the polarity of both wildtype (Genotype 2 and Genotype 4) and *stan^-^* receiving cells (Genotype 3 and Genotype 5) to the same extent. They also affect *pk-sple^-^ stan^+^* (Genotype 17 and Genotype 18) and *pk-sple^-^ stan^-^* (Genotype 19 and Genotype 20) receiving cells with the same strength (Figure 3 and Figure S3). These results show that neither Pk, Sple nor Stan are required for polarity signalling by the Ds/Ft system, although Pk and Sple can change the sign of the response; suggesting that the formation of Ds-Ft bridges works well even the absence of Pk/Sple and/or the Stan system. Any effect seen has to be downstream of these bridges. They also show that Pk and Sple do not act as an essential link between the Ds/Ft system and the Stan system, because if they were such a link, removal of Pk and Sple would block effects on polarity caused by overexpressing *ft*.

Results supporting these conclusions came from clones that altered the Ds/Ft system in flies whose polarity was altered by excess Pk or Sple:

2. In flies in which *pk* or *sple* are overexpressed. Clones that lack *ft* made in flies in which *pk* is generally overexpressed (Genotype 21) behave as follows: where the polarity of much of the surrounding background is reversed from normal, with the hairs pointing forwards (ie in the A compartment), *ft^-^* clones act with the opposite sign to that in the wildtype (Genotype 22) and hairs around the clone tend to point outwards (Figure 6). In the P compartment, where overexpression of *pk* produces no change to polarity, the *ft^-^* clones behave as they do in the wildtype, that is the hairs point outwards from the clone (Figure 6).

**Figure 5.**
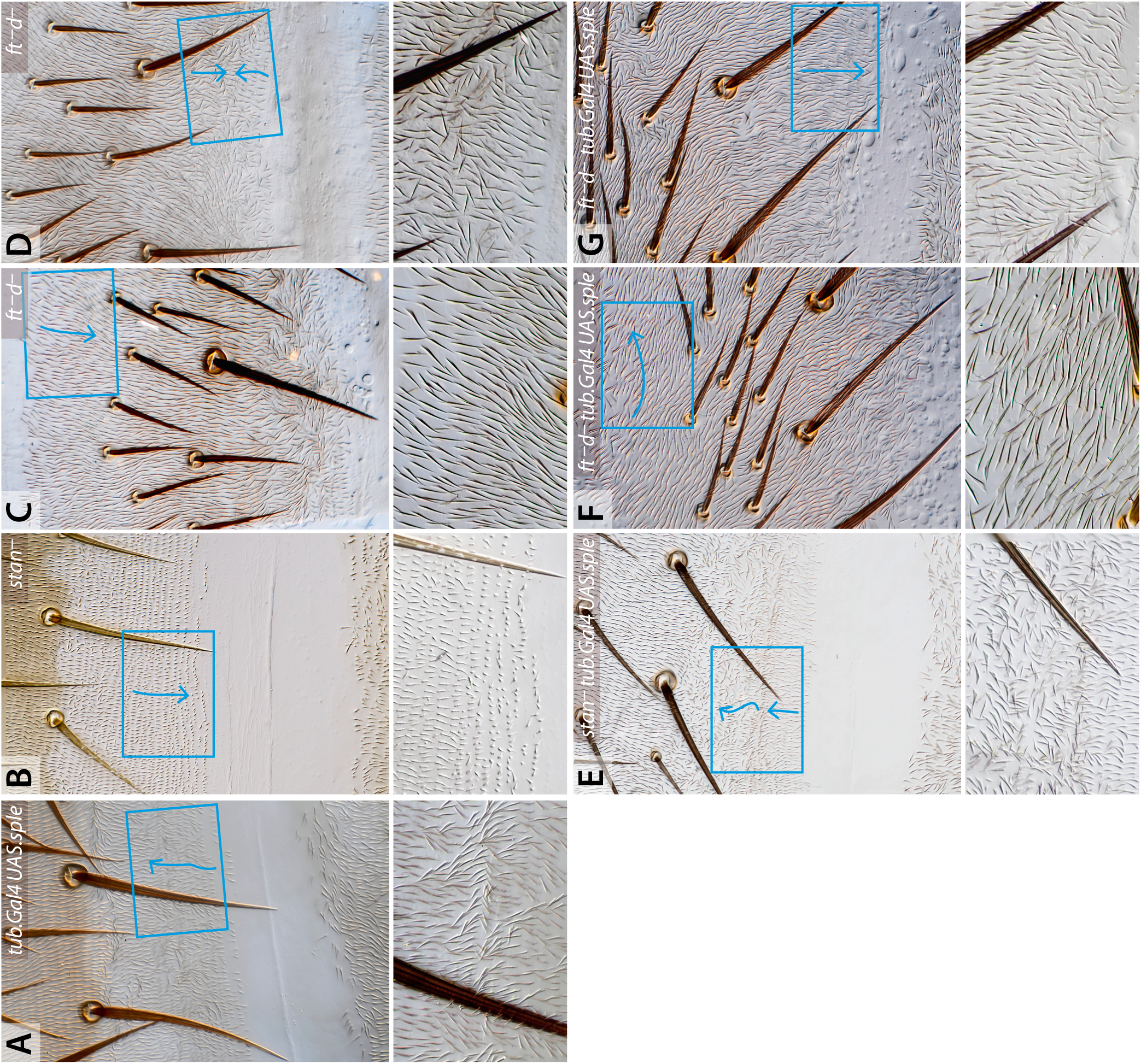
Effects of overexpressing *sple* on polarity of cells in which either the Stan system (*stan^-^*) or the Ds/Ft system is broken (*ft^-^ d^-^*). Overexpression of *sple* in *stan^-^* and the wildtype reverses all or most of the P compartment to point forwards (compare A, B and E) but overexpression of *sple* in a *ft^-^ d^-^* background produces a P compartment of normal polarity (G) and even the rear of the P region, which points forward in *ft^-^ d^-^* (D) is now “rescued” to normal polarity. Overexpression of *sple* in *ft^-^ d^-^* flies also alters the polarity at the front of the A compartment (C and F) turning the hairs laterally, while overexpressing *pk* turns the hairs to point anteriorly (Figure 4). Compare Figure S4 for expression of *sple* in *d^-^* flies

**Figure 6.**
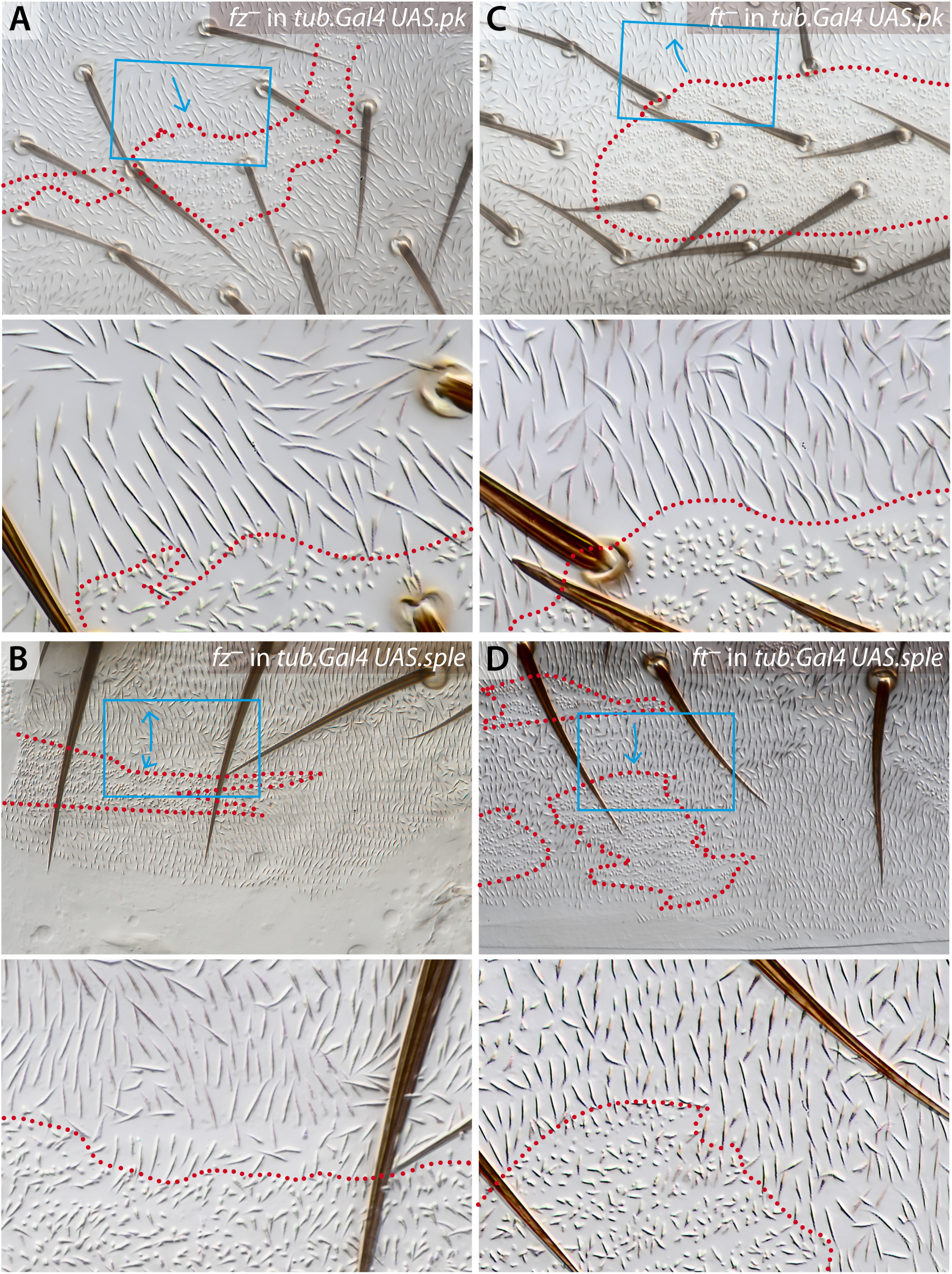
Behaviour of *fz^-^* and *ft^-^* clones in flies overexpressing isoforms of the *pk* gene. *fz^-^* clones behave normally, polarising receiving cells inwards in both A and P either in *tub.Gal4 UAS.pk* or *tub.Gal4 UAS.sple* flies, independently of the polarity of their surrounds (A and B). The effects of *ft^-^* clones, but only in territories with reversed polarity, are the opposite of normal: in the wildtype these effects are inwards in A, outwards in P while in *tub.Gal4 UAS.pk* the cells close to the anterior clones point outwards (C) and in *tub.Gal4 UAS.sple* the cells nearby the posterior clones point inwards (D). See Figure S5 for analysis of maximum range of effects of *fz^-^* clones.

Clones that lack *ft* made in flies in which *sple* is generally overexpressed (Genotype 23) behave as follows: in the A compartment, which has normal polarity, these clones affect these receiving cells as they affect wildtype cells; hairs around the clone point inwards (Figure 6). In the P compartment, where the polarity of the surrounding background is reversed from normal with the hairs pointing forwards, the *ft^-^* clones now polarise receiving cells with the opposite sign to that in the wildtype, that is the hairs point inwards into the clone (Figure 6).

In the A compartment of the abdomen, clones that lack *ds* have effects of the opposite sign to *ft^-^* clones in both classes (see previous points 1 and 2) of experiments above —as would be expected. However *ds^-^* clones have little or no effect in the P compartment in all genotypes tested (data not shown, Genotype 24, Genotype 25 and Genotype 26).

3. These results show that the Ds/Ft system can function independently of Pk and Sple but that Pk and Sple can modulate the sign of its output. This dramatic effect could, in principle, be due to Pk and/or Sple affecting the patterns of expression of *ds*, and/or *fj* and thereby changing the orientation of the Ds/Ft system gradients. To test we studied the expression of enhancer traps for *ds* and *fj* loci in *pk-sple^-^* flies and saw no departure from the wildtype patterns (Genotype 28, Genotype 29, Genotype 30 and Genotype 31; Figure S6). It follows that Pk and Sple determine whether polarised structures in the cell, the hairs and bristles, point up or down the gradients of Ds and Fj.

#### (iv) Pk and Sple alter polarity even when the Stan system is broken

Were Pk/Sple to act only upstream of the Stan system one would not expect overexpression of *pk* or *sple* to have an effect on the polarity of *stan^-^*flies. However uniform overexpression of *pk* causes large changes of polarity in the abdomen of wildtype flies and flies with a broken Stan system (Genotype 32) in the A compartment, without affecting the P compartment (Figure 4). While generalised overexpression of *sple* also affects the polarity of both wildtype and *stan^-^* flies — altering the polarity of the P compartment of the abdomen, without much affecting the A compartment (Genotype 33, Figure 5).

#### (v) Pk and Sple affect PCP even when the Ds/Ft system is broken

If Pk and Sple acted exclusively on the Ds/Ft system, one would not expect Pk and Sple proteins to affect PCP if the Ds/Ft system were broken.

1. General overexpression of *pk* or *sple* in a broken Ds/Ft system. But we find that ubiquitous overexpression of *pk* alters polarity of the A compartment (and part of the P compartment) of *ds^-^pk-sple^-^* (Genotype 34, Figure 1), *d^-^* (Genotype 35, Figure S4) and *ft^-^ d^-^* flies (Genotype 36, Figure 4). Similarly, general overexpression of *sple* affects the polarity of the P compartment of the abdomen of *ds^-^pk-sple^-^* (Genotype 37, Figure 1), *d^-^* (Genotype 38, Figure S4) and *ft^-^ d^-^* flies (Genotype 39, Figure 5).

In *d^-^* flies, the A and P compartments are largely normal but a section of the P compartment is reversed, as in *ds^-^* (or *ft*) flies. When ubiquitous Pk is added to *d^-^* or *ft^-^ d^-^* flies, the anterior part of the A compartment is altered to point forwards and the reversed rear section of the P compartment is “rescued” so that it points backwards, as in the wildtype. Thus Pk affects both the A and the P compartment in these flies. However, unlike Pk, ubiquitous Sple affects *d^-^* and *ft^-^ d^-^* flies differentially: in a *d^-^* background there is no change to the A compartment, but the whole P compartment is largely reversed. But, in a *ft^-^ d^-^* background the anterior region of the A compartment points laterally and, as noted by Sharp and Axelrod (2016) the P compartment is rescued, having a normal orientation — thus Pk and Sple have similar effects on *ft^-^ d^-^* but very different effects on *d^-^* flies. It follows from these findings that Ft has outputs that are independent of D and that these outputs are altered by Sple but not by Pk. Note that both Sple and Pk can rescue the reversed polarity in the P compartment in a completely broken Ds/Ft system (*ft^-^ d*^-^) perhaps through their effects on the Stan system or, maybe, through other contributors to PCP (Figure 4, Figure 5 and Figure S4).

2. Clones that overexpress *pk* or *sple* in a broken Ds/Ft system. We find that clones of cells overexpressing *sple* (Genotype 40; Lawrence et al., 2004) or *pk* (Genotype 41; data not shown), have small non-autonomous effects in the wildtype and, more so, in *ds^-^* flies (Genotype 42 and Genotype 43) where they polarise receiving cells to point strongly inwards (Figure 7). Perhaps these clones act via the Stan system? It is pertinent that both wing and abdominal cells that overexpress the *pk* gene accumulate Vang uniformly on the cell membrane (Bastock et al., 2003; Olofsson et al., 2014). If this were to happen in our experiments, then the clone could behave as if it were overexpressing *Vang* and should polarise surrounding cells inwards, as observed; this effect should be stronger in *ds^-^* than in *ds^+^* cells, also as observed. To test this hypothesis further we made *Vang* clones that overexpressed *pk* (Genotype 44), as well as control *Vang* clones (Genotype 45), in *ds^-^* flies. Both these types of clones behaved like *Vang* clones in wildtype flies (Genotype 46), and could not be distinguished from each other, ie they polarise *ds^-^* receiving cells strongly outwards (Figure 7), confirming the hypothesis that cells overexpressing *pk* polarise cells because they accumulate Vang, a Stan system protein. Thus, overexpressing Pk interferes with the Stan system. These results show that Pk and Sple do have functions that are independent of the Ds/Ft system.

**Figure 7.**
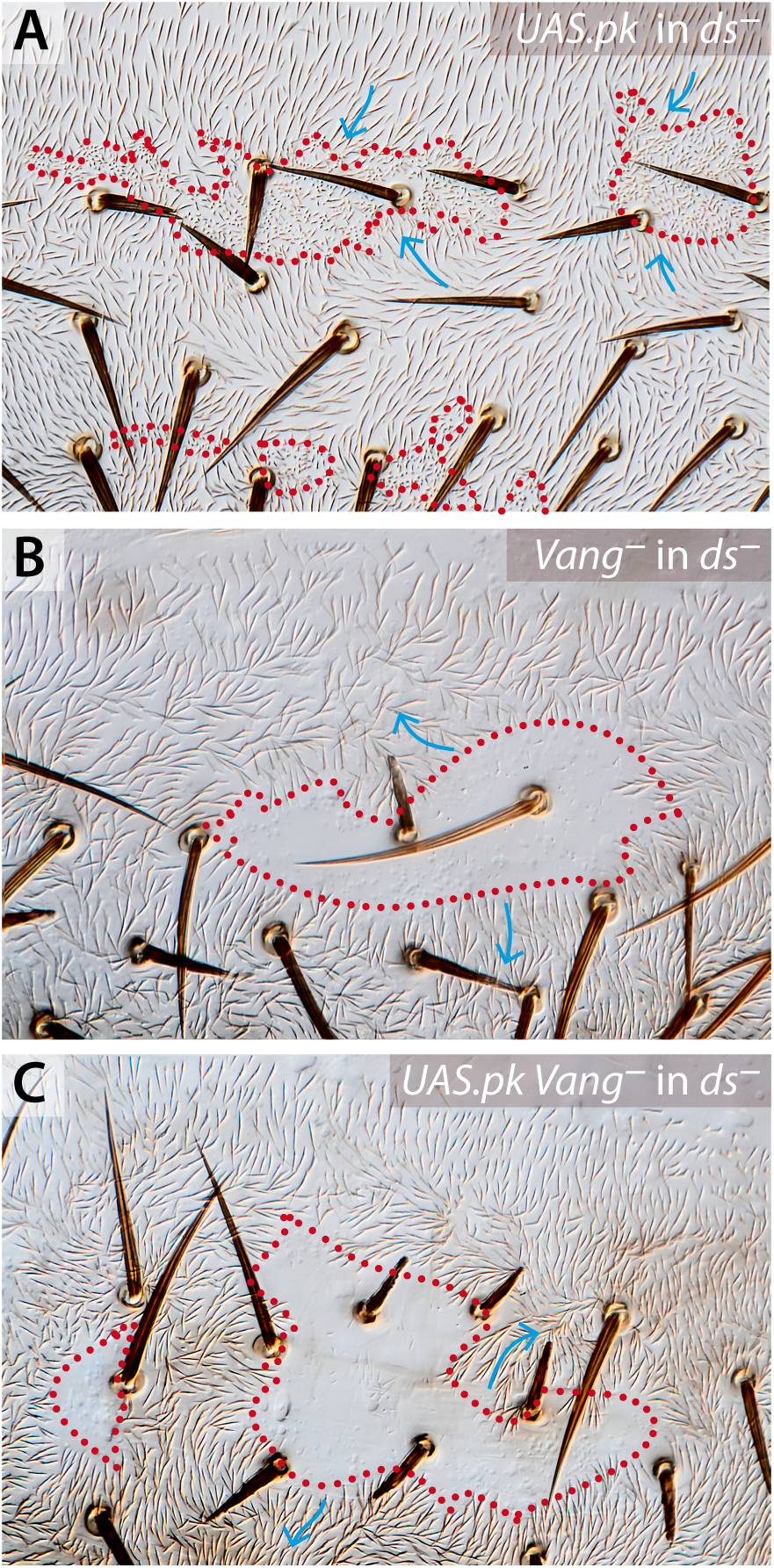
Effects of pk-expressing clones in flies broken for the Ds/Ft system. Clones that overexpress *pk* polarise *ds^-^* cells strongly inwards (A). Clones lacking *Vang* (B) as well as clones that, lacking *Vang*, also overexpress *pk* (C), polarise *ds^-^* receiving cells strongly outwards.

#### (vi) Can Pk function independently of both systems of PCP?

We have presented evidence that the Pk and Sple modulate, in different ways, both the Ds/Ft and the Stan systems. We wondered if Pk and Sple proteins could have outputs independently of these systems. To test, we have overexpressed *pk* in a genetic background in which both systems are broken, viz: *ft^-^ d^-^ stan^-^*. We found, using blind screening, that could not distinguish the phenotypes of these flies with and without the overexpression of *pk* (Genotype 47 and Genotype 48, Figure 4), arguing that the polarity effects of an excess of Pk are due to Pk acting through the two systems and by no other route.

## Discussion

Our aim is to understand the contribution of Pk and Sple to building planar cell polarity in the wildtype fly. The main results and conclusions are listed below.

### The Ds/Ft system and the Stan system act independently and are not linked via Sple and/or Pk

*ft*-overexpressing clones reorient wildtype receiving cells, outwards in the A compartment (Casal et al., 2006) and inwards in the P (this paper). These clones have the same effects on cells in which the Stan system of PCP is broken (for example in *stan^-^* flies; Figure 2, Figure 3 and Figure S3). It follows that this reorientation of polarity cannot be due to any intracellular interaction between Stan and any component of the Ds/Ft system within the sending cells. However it could be argued that extra Ft in the sending cell, attracting Ds in the receiving cell, would, non-autonomously, influence some residual capability of the Stan system in the receiving *stan^3^/stan^E59^* cells to respond and propagate polarity to neighbouring cells. Yet, even clones that overexpress both *fz* and *stan* (ie cells that have a fully functional Stan system) fail to repolarise *stan^3^/stan^E59^* cells (Casal et al., 2006). Thus the propagation of a polarity change observed around cells that overexpress *ft* cannot be due to any non-autonomous effect on the Stan system. These results show that, at least for both compartments of the abdomen, the Ds/Ft system acts independently of the Stan system (Casal et al., 2006; Lawrence et al., 2007; Lawrence, 2011).

Here we make *stan^E59^* clones (*stan^E59^* introduces a premature stop codon in the ectodomain Usui et al., 1999) that overexpress *ft* or *ds* in *pk-sple^-^ stan^3^/ pk-sple^-^ stan^E59^* flies; and show that these clones repolarise the receiving cells. This polarisation cannot depend on Pk and Sple intervening, inside the cells of the clone, between the Ds/Ft and the Stan systems because the sending cells lack the *stan* and *pk* genes completely; in addition the receiving cells lack the *pk* gene and any functional Stan (see previous paragraph). A finding that conflicts with current models in which the *pk* gene products are proposed to link the two systems of PCP (Hogan et al., 2011; Ayukawa et al., 2014; Merkel et al., 2014; Olofsson et al., 2014; Ambegaonkar and Irvine, 2015).

Another argument is relevant here: clones affecting the Ds/Ft system have outputs of different sign in the A and P compartments (Casal et al., 2002). If this divergent polarisation were to act through and depend on the Stan system via a molecular link of Pk and/or Sple, then we would expect the polarising output from Stan system clones (eg from *fz^-^* clones) to be also of different sign in the two compartments and to be dependent on that link (cf Figure 7 in Ayukawa et al., 2014). However, this is not the case (Figure S2E and Figure S2F).

### Pk/Sple act independently of the Stan system

Loss of the *pk* gene or overexpressing the Pk isoform reverses polarity of most of the A compartment, having strong effects even in flies with a broken Stan system (*stan^-^*) Similarly, overexpressing Sple reverses polarity in the P compartment in *stan^-^* flies; it follows that Pk and Sple can act independently of the Stan system. This does not fit easily with the current view that Pk functions as an essential component of the Stan system; for example the lack of requirement for the *pk* gene contrasts with a strong requirement for the other key Stan system genes (ie Fz, Stan or Vang) in the receiving cells (Taylor et al., 1998; Lawrence et al., 2004; Strutt and Strutt, 2007; Strutt and Warrington, 2008; Struhl et al., 2012).

The Stan system proteins Stan, Fz, Vang and Pk are all preferentially localised to specific regions of the cell membrane and this is considered to be important for their functions in PCP. Nevertheless, *pk-sple^-^* receiving cells, in which Stan, Fz and Vang no longer appear to be localised (reviewed in Strutt, 2009), can respond at least as well to such sending cells as wildtype ones (Adler et al., 2000; Lawrence et al., 2004). This dilemma could be resolved if the observed asymmetry were not so directly related to function as has been assumed and were more a consequence than a cause of polarity (Lawrence et al., 2004).

### Pk and Sple modulate the Ds/Ft system, determining the polarity of its output

Sending cells that overexpress *ds* or *ft*, or lack *ds* or *ft*, change the polarity of receiving cells, even in the absence of Pk and Sple— it follows that these proteins cannot be necessary for the Ds/Ft system to function and propagate polarity from cell to cell. However the sign of this change depends on whether the receiving cells contain, lack or overexpress products of the *pk* gene. These results show that the Pk and Sple can alter the sign of polarisation that is produced by the Ds/Ft system. But, how do Pk and Sple have their various effects on polarity? It appears that the sign of polarisation depends on the relative amounts of Pk and Sple in a particular region of the fly (Gubb et al., 1999; Ayukawa et al., 2014). One model is that the Ds/Ft proteins might act through Pk and Sple to bias the orientation of microtubules and thus PCP —because oriented microtubules could transport Stan system components preferentially to one side of the cell (Shimada et al., 2006; Harumoto et al., 2010; Matis et al., 2014; Olofsson et al., 2014). But, correlation between microtubule orientation, Pk/Sple function, and PCP is inconsistent or lacking, clearly so in the distal half of the wing and the P compartment of the adult abdomen (Harumoto et al., 2010; Sharp and Axelrod, 2016), leading to doubts about the validity of the hypothesis (Ambegaonkar and Irvine, 2015). Also, this model is now contradicted by our results, which show that abnormal amounts of Ds, Ft, Sple or Pk can all affect PCP even when the Stan system is broken.

A diagram suggesting how the *pk* gene might fit into the organisation of PCP is given in Figure 8.

**Figure 8.**
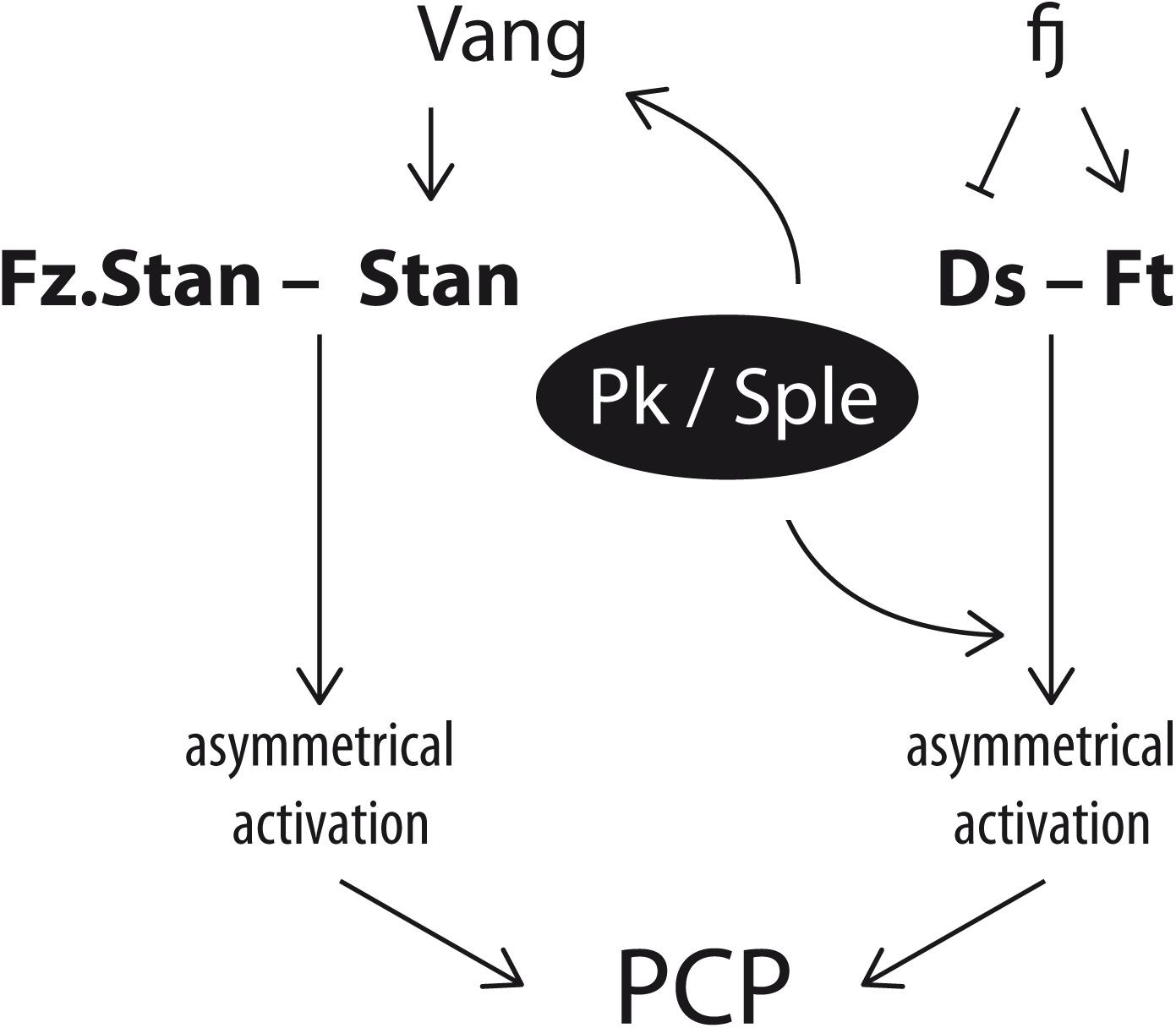
Pk and Sple functions in the context of PCP. PCP depends on molecular bridges between cells: for the Stan system the key bridge consists of a complex of Stan and Fz in one cell and Stan in the other; Vang promotes function of the Stan pillar of this bridge (Struhl et al., 2012). For the Ds/Ft system, Ds in one cell is linked to Ft in another, the activity of both is modulated by Fj (reviewed in Butler and Wallingford, 2017). Pk and/or Sple bind to Vang and promote asymmetrical distribution of Vang and other PCP molecules. Yet in the absence of Pk and Sple, the Stan system can still receive and send polarity information, implying that it is the asymmetric activation of protein complexes that polarise a cell rather than asymmetric localisation. Pk and Sple alter the sign of the polarity output of the Ds/Ft system, but by an unknown mechanism. Yet, Pk and Sple can alter polarity output even when the Ds/Ft system is broken. The results show that Pk and Sple can act separately on both systems, implying some general function of Pk and Sple in cell polarity. The indispensable elements of the two systems are shown in bold.

### The functions of Pk and Sple

It has been suggested that Pk and Sple do fundamentally different things (Ayukawa et al., 2014; Ambegaonkar and Irvine, 2015); however our findings fit better with the view that the two isoforms have similar molecular functions and the differences between them are due to their expression in different patterns (Gubb et al., 1999). Indeed, in the results (section **v**) where we study the behaviour of *ft^-^* or *ft*-expressing clones, we found that removal of Pk and Sple or ubiquitous expression of either can eliminate any differences in responses between the A and the P compartment cells.

It might appear that ectopic Pk can act in only the A compartment and Sple in the P, but it cannot be so simple, for the universal expression of either Pk or Sple can rescue the reversed polarity at the back of the P compartment in *ft^-^ d^-^* flies (Figure 4, and Sharp and Axelrod, 2016). Also, when *sple* is generally overexpressed in *ft^-^ d^-^* flies, polarity of the anterior region of the A compartment is considerably altered (Figure 5). Looking at the P compartment, the action of Pk appears to be independent of an intact Ds/Ft system, but the effects of Sple in the P compartment depend on whether the background genotype is *d^-^* or *ft^-^ d^-^* (Figure S4). Part of this difference could be due to direct interaction between Sple and Ds but not between Pk and Ds (Ambegaonkar and Irvine, 2015). However Ayukawa et al. (2014) find that both Sple and Pk bind to each other and to D (but not to Ds), suggesting the difference may have other causes.

But why are the Pk and Sple proteins asymmetrically localised in the cell? Part of the answer could be that Pk and Sple work with and/or bind to components of the Ds/Ft system which are themselves asymmetrically localised. (Ayukawa et al., 2014; Ambegaonkar and Irvine, 2015). But this cannot be all of the answer as Pk is not properly localised in *stan^-^* cells (Tree et al., 2002b), in which Ds and Ft are, presumably, normally localised.

How can we understand the effect of Pk and Sple on the Stan system, particularly on range? In the A compartment, a high level of Sple reduces polarity changes induced by *fz^-^* clones, while the loss of the *pk* gene increases their range. One explanation could depend on Sple and Pk (or the lack of these proteins) acting on the Ds/Ft system —if they made the polarity induced by Ds/Ft in the cells more (or less) robust it would make it more difficult (or easier) for clones affecting the Stan system to alter PCP. Another explanation could relate to some direct effect of Pk (and Sple) on Vang (Bastock et al., 2003) which fits our observations with clones overexpressing Pk (Figure 7). The function of Vang in the Stan system is somewhat unclear; like Pk, Vang is present in larger than stoichiometric amounts in relation to the two molecules that form the intercellular bridge, Stan and Fz (Strutt et al., 2016), yet affects bridge function (Struhl et al., 2012). The abdominal phenotypes of *Vang^-^* and *pk-sple^-^* are somewhat similar, both having areas of reversed polarity (Lawrence et al., 2004), suggesting a commonality of function. Indeed there is a recent model proposing that Pk acts on the stability of Fz intracellularly (via Dsh) and in the adjacent cell (via Vang); the former effect may involve endocytosis (Warrington et al., 2017). Our experiments argue that the function of Pk is not limited to the Stan system but includes, independently, the Ds/Ft system. In any case we have no explanation for the lack of apparent effects of Pk and Sple on the range of *fz^-^* clones in the P compartment.

What could be the purpose of such complexity? In *Drosophila* the consistent orientation of the wing hairs may have led to an oversimplified and idealised picture. Elsewhere, the presentation of PCP is more complex: consider the mixed orientation of rows of hairs and denticles on the Drosophila larva, differing dorsally and ventrally, or, in mammals, the startlingly diverse orientation of stereocilia in the vestibular system, or the complex patterns of hair orientation on the skin. Two separate genetic systems which generate polarity by reading the slopes of morphogen gradients, plus Pk and Sple to modulate output in different parts of the body, could generate much of this flexibility in PCP.

## Conclusion

We have found for the abdomen that Pk and Sple are not essential components of either the Ds/Ft or the Stan systems. We have shown that they do not function as a link between the two systems. Instead, Pk and Sple appear to modulate the polarity outputs of both the Ds/Ft system and the Stan system with the most conspicuous effects on the former. Both these systems differ in their components but are similar in their logic; both utilise intercellular molecular bridges that become distributed asymmetrically within each cell. Pk and Sple could help produce this asymmetry— perhaps via a generic function in cell biology whose mechanism is still undescribed. Our genetic experiments on the abdomen point to conclusions that differ from the prevailing view that the *pk* gene mediates between the Ds/Ft and the Stan systems. We do not know if our conclusions apply to other organs in the fly, but if we adopt the hypothesis that they do, they suggest that current views of the wildtype functions of the *pk* gene should be reconsidered.

## Materials and Methods

### Mutations and transgenes

The FlyBase (Gramates et al., 2017) entries for relevant mutations and transgenes are the following: *tub.Gal4: Scer\GAL4^alphaTub84B.PL^. tub.Gal80: Scer\GAL80^alphaTub84B.P^*. *UAS.ectoDs: ds^ecto.Scer\UAS^*. *UAS.ft: ft^Scer\UAS.cMa^. UAS.fz : fz^Scer\UAS.cSa^. UAS.pk: pk^Scer\UAS.cGa^. UAS.sple: pk^sple.Scer\UAS^. ck^UAH21^. d^GC13^. ds^UA071^* and *ds^2D60b^. fj^p1^. ft^8^* and *ft^G-rv^. fz^15^. pk^pk-sple-13^. pwn^1^. sha^1^. stan^3^* and *stan^E59^. trc^1^*.

### Experimental Genotypes

Genotype 1: ***UAS.fz* clones in *stan^-^* flies:** *y w hs.FLP; FRT42D tub.Gal80 stan^3^ hs.CD2, y^+^/FRT42D pwn stan^E59^; UAS.fz/ tub.Gal4*
Genotype 2: ***UAS.ft* clones in wild type flies:** *y w hs.FLP tub.Gal4 UAS.nls-GFP/y w hs.FLP; d^GC13^ FRT42D pwn sha/d^GC13^ FRT42D tub.Gal80, y^+^; UAS.ft/+*
Genotype 3: ***UAS.ft* clones in *stan^-^* flies:** *y w hs.FLP; FRT42D pwn stan^E59^ sha/FRT42D tub.Gal80 stan^3^ hs.CD2, y^+^; UAS.ft/ tub.Gal4*
Genotype 4: ***UAS.ectoDs* clones in wild type flies:** *y w hs.FLP tub.Gal4 UAS.nls-GFP/ y w hs.FLP; FRT42D pwn stan^E59^ sha/ FRT42D tub.Gal80; UAS.ectoDs/ +*
Genotype 5: ***UAS.ectoDs* clones in *stan^-^* flies:** *y w hs.FLP; FRT42D pwn stan^E59^ sha/FRT42D tub.Gal80 stan^3^ hs.CD2y^+^; UAS.ectoDs/ tub.Gal4*
Genotype 6: ***UAS.fz* clones in wild type flies:** *y w hs.FLP; FRT42D pwn/ FRt42D tub.G80, y^+^; tub.Gal4/ UAS.fz*
Genotype 7: ***UAS.fz* clones in *ds^-^* flies:** *y w hs.FLP tub.Gal4 UAS.nls-GFP/y w hs.FLP; ds^UA071^ FRT42D pwn/ ds^UA071^ FRT42D tub.Gal80; UAS.fz hs.CD2, y^+^/ +*
Genotype 8: ***ds^-^ pk^-^ sple^-^* flies:** *y w hs.FLP; ds^UA071^ pk^pk-sple–13^; UAS.sple/ TM2*
Genotype 9: ***pk-sple^-^ stan^-^* flies:** *y w hs.FLP tub.Gal4 UAS.nls-GFP; FRT42D pk^pk-sple-13^ stan^3^ tub.Gal80/ FRT42D pk^pk-sple–13^ stan^E59^ sha; MRS/ TM2*
Genotype 10: ***stan^-^* flies:** *y w hs.FLP; FRT42D tub.Gal80 stan^3^ hs.CD2, y^+^/ FRT42D pwn stan^E59^; MRS/ TM2*
Genotype 11: ***pk-sple^-^* flies:** *pk^pk-sple–13^*
Genotype 12: ***UAS.fz* clones in *pk^-^ sple^-^* flies:** *y w hs.FLP tub.Gal4 UAS.nls-GFP/y w hs.FLP; FRT42D pk^pk-sple–13^ sha/ FRT42D pk^pk-sple–13^ tub.Gal80; UAS.fz fz^15^ fz2^C1^ FRT2A/ +*
Genotype 13: ***fz^-^* clones in *pk-sple^-^* flies:** *y w hs.FLP122; FRT42 pk^pk-sple–13^/ CyO; fz[P21] trc FRT2A/ tub.Gal80 FRT2A*
Genotype 14: ***fz^-^* clones in *tub.Gal4 UAS.sple* flies:** *y w hs.FLP tub.Gal4 UAS.nls-GFP/ w; UAS.sple/ +; UAS.sple; fz^15^ trc^C1^ FRT2A / hs.GFPw^+^ hs.CD2, y^+^ ri FRT2A*
Genotype 15: ***fz^-^* clones in *tub.Gal4 UAS.pk* flies:** *y w hs.FLP tub.Gal4 UAS.nls-GFP/ w; UAS.pk/ +; UAS.sple; fz^15^ trc^C1^ FRT2A / hs.GFPw^+^ hs.CD2, y^+^ ri FRT2A*
Genotype 16: ***fz^-^* clones in *ds^-^* flies:** *ds^UA071^ FRT39/ ds^33k^ bw^V1^; fz^H51^ trc^C1^ ri FRT2A/ hs.CD2, y^+^ hs.GFP ri FRT2A/ TM3*
Genotype 17: ***UAS.ft* clones in *pk****^-^* ***sple^-^* flies:** *y w hs.FLP122 tub.gal4 UAS.nls-GFP/y w hs.FLP; FRT42D pk^pk-sple-13^ sha/ FRT42 pk^pk-sple-13^ tub.Gal80/ UAS.ft/ +*
Genotype 18: ***UAS.ectoDs* clones in *pksple^-^* flies:** *y w hs.FLP122 tub.gal4 UAS.nls-GFP/ y w hs.FLP; FRT42D pk^pk-sple-13^ sha/ FRT42 pk^pk-sple-13^ tub.Gal80/ UAS.ectoDs/ +*
Genotype 19: ***UAS.ectoDs* clones in *pk****^-^****sple^-^ stan^-^* flies:** *y w hs.FLP tub.Gal4 UAS.nls-GFP/y w hs.FLP; FRT42D pk^pk-sple-13^ stan^3^ tub.Gal80, y^+^/FRT42 pk^pk-sple-13^ stan^E59^ sha ; UAS.ectoDs/ +*
Genotype 20: ***UAS.ft* clones in *pk****^-^* ***sple^-^ stan^-^* flies:** *y w hs.FLP tub.Gal4 UAS.nls-GFP/ y w hs.FLP122; FRT42D pk^pk-sple-13^ stan^3^ tub.Gal80, y^+^/FRT42D pk^pk-sple-13^ stan^E59^ sha ; UAS.ft/ +*
Genotype 21: ***ft^-^* clones in *tub.Gal4 UAS.pk* flies:** *y w hs.FLP tub.Gal4 UAS.GFP-nls/ y; ft^15^ stc FRT39/ FRT39; UAS.pk/ +*
Genotype 22: ***ft^-^* clones in wild type flies:** *y w hs.FLP; ft^15^ stc FRT39/ FRT39*
Genotype 23: ***ft^-^* clones in *tub.Gal4 UAS.sple* flies:** *y w hs.FLP tub.Gal4 UAS.GFP-nls/ y; ft^15^ stc FRT39/ FRT39; UAS.sple/ +*
Genotype 24: ***ds^-^* clones in *tub.Gal4 UAS.sple* flies:** *w hs.FLP tub.Gal4 UAS.nls-GFP; ds^UA071^ ck^UAh21^ FRT40A/ FRT40A; UAS.sple/ +*
Genotype 25: ***ds^-^* clones in *tub.Gal4 UAS.pk* flies:** *w hs.FLP tub.Gal4 UAS.nls-GFP; ds^UA071^ ck^UAh21^ FRT40A/ FRT40A; UAS.pk/ +*
Genotype 26: ***ds^-^* clones in wild type flies:** *y w hs.FLP/* +; *ds^UA071^ ck^UAh21^ FRT40A/ Dp*(*1;2*)*sc^19^ w^+30c^ FRT40A*
Genotype 27: **Control clones in *pk^-^sple^-^ stan^-^* flies:** *y w hs.FLP tub.Gal4 UAS.nls-GFP/ y w hs.FLP; FRT42D pk^pk-sple-13^ stan^E59^ sha/ FRT42D pk^pk-sple-13^ stan^3^ tub.Gal80, y^+^; MRS/ +*
Genotype 28: ***ds.lacZ* flies:** *y hs.FLP/* +; *ds^2D60b^ FRT42D pk^pk-sple-13^/ +*
Genotype 29: ***fj.lacZ* flies:** *y hs.FLP/* +; *FRT42D pk^pk-sple-13^ fj^P1^/ +*
Genotype 30: ***ds.lacZ pk-sple^-^* flies:** *y hs.FLP/* +; *ds^2D60b^ FRT42D pk^pk-sple-13^/ FRT42D pk^pk-sple-13^*
Genotype 31: ***pk-sple^-^ fj.lacZ* flies:** *y hs.FLP/* +; *FRT42D pk^pk-spk 13^ fj^P1^/ FRT42D FRT42D pk^pk-sple-13^*
Genotype 32: ***stan^-^ tub.Gal4 UAS.pk* flies:** *y w hs.FLP tub.Gal4 UAS.nls-GFP/y w hs.FLP; FRT42D pwn stan^E59^ sha/ FRT42D stan^3^; UAS.pk/ TM2*
Genotype 33: ***stan^-^ tub.Gal4 UAS.sple* flies:** *y w hs.FLP tub.Gal4 UAS.nls-GFP/y w hs.FLP; FRT42D pwn stan^E59^ sha/ FRT42D stan^3^; UAS.sple/ TM2*
Genotype 34: ***ds^-^pk^-^sple^-^ tub.Gal4 UAS.pk* flies:** *y w hs.FLP tub.Gal4 UAS.nls-GFP/ y w hs.FLP; ds^UA071^ pk^pk-sple-13^; UAS.pk/ TM2*
Genotype 35: ***d^-^ tub.Gal4 UAS.pk* flies:** *y w hs.FLP tub.Gal4 UAS.nls-GFP/y w hs.FLP; d^GC13^ pr cn/ft^G-rv^ d^GC13^ FRT40; UAS.pk/ +*
Genotype 36: ***ft^-^ d^-^ tub.Gal4 UAS.pk* flies:** *y w hs.FLP tub.Gal4 UAS.nls-GFP/y w hs.FLP; ft^8^ d^GC13^ FRT40A/ft^G-rv^ d^GC13^ FRT40A; UAS.sple/ +*
Genotype 37: ***ds^-^pk^-^ sple^-^ tub.Gal4 UAS.sple* flies:** *y w hs.FLP tub.Gal4 UAS.nls-GFP/ y w hs.FLP; ds^UA071^ pk^pk-sple-13^; UAS.sple/ TM2*
Genotype 38: ***d^-^ tub.Gal4 UAS.sple* flies:** *y w hs.FLP tub.Gal4 UAS.nls-GFP/ w; ft^G-rv^ d^GC13^/ d^GC13^ pr cn; UAS.sple/ +*
Genotype 39: ***ft^-^ d^-^ tub.Gal4 UAS.sple* flies:** *y w hs.FLP tub.Gal4 UAS.nls-GFP/y w hs.FLP; ft^8^ d^GC13^ FRT40A/ft^G-rv^ d^GC13^ FRT40A; UAS.sple/ +*
Genotype 40: ***UAS.sple* clones in wild type flies:** *y w hs.FLP tub.Gal4 UAS.nls-GFP/ y w hs.FLP;FRT42D pwn/FRT42D tub.Gal80; UAS.sple/ +*
Genotype 41: ***UAS.pk* clones in wild type flies:** *y w hs.FLP tub.Gal4 UAS.nls-GFP/y w hs.FLP;FRT42D pwn/ FRT42D tub.Gal80; UAS.pk/ +*
Genotype 42: ***UAS.sple* clones in *ds^-^* flies:** *y w hs.FLP tub.Gal4 UAS.nls-GFP/y w hs.FLP; ds^UA071^ ck^UAh21^ FRT40A/ ds^UA071^ tub.Gal80 FRT40A; UAS.sple/ MRS*
Genotype 43: ***UAS.pk* clones in *ds* flies:** *y w hs.FLP tub.Gal4 UAS.nls-GFP/y w hs.FLP; ds^UA071^ ck^UAh21^ FRT40A/ ds^UA071^ tub.Gal80 FRT40A; UAS.pk/MRS*
Genotype 44: ***Vang^-^ UAS.pk* clones in *ds^-^ flies: y w hs.FLP tub.Gal4 UAS.nls-GFP/y w hs.FLP; ds^UA071^ FRT42D tub.Gal80/ ds^UA071^ hs.CD2, y^+^*** *FRT42D pwn Vang^tbm-6^ sha; UAS.pk/ +*
Genotype 45: ***Vang^-^* clones in *ds^-^* flies:** *y w hs.FLP; ds^UA071^ FRT42D tub.Gal80/ ds^UA071^ hs.CD2, y^+^ FRT42D pwn Vang^stbm-6^ sha; UAS.pk/ +*
Genotype 46: ***Vang* clones in wild type flies:** *y/y hs.FLP; FRT42D pwn Vang^stbm-6^ FRT42D hs.CD2, y^+^*
Genotype 47: ***ft^-^ d^-^ stan^-^ tub.Gal4 UAS.pk* flies:** *y w hs.FLP tub.Gal4 UAS.nls-GFP/y w hs.FLP ; ft^Gv-r^ d^GC13^ stan^3^/ ft^8^ d^GC13^ stan^E59^; TM2/ UAS.pk*
Genotype 48: ***ft^-^ d^-^ stan^-^* flies:** *y w hs.FLP ; ft^Gv-r^ d^GC13^ stan^3^/ft^8^ d^GC13^ stan^E59^; TM2/ UAS.pk*

### Clone induction and microscopy

Clones were induced by heat shocking third instar larvae for 1 hr at 34°C. Adult abdominal cuticles were studied as before (e.g., Lawrence et al., 2004; Casal et al., 2006).

### Quantitation

Individual hairs along the entire perimeter of each clone (about 60–100 hairs per clone) were each scored as pointing largely into, outwards or parallel to the clone. Parallel hairs, which averaged 8% of the hairs, were counted; half was added equally to the inwards and outwards sets. The average orientation is then found for each clone (between 10 and 20 clones per genotype).

For range measurments, for each clone (n=20) the maximum extent in cell rows of the induced polarity changes was measured. The observer was blinded as to genotypes; he chose clones located in the middle of the A compartment and the middle or rear of the hairy region of the P compartment; small clones were avoided. Statistical analysis and graphics were performed in R using standard packages (R Core Team, 2016) and the *reshape* and *ggplot* packages (Wickham, 2007; Wickham, 2009).

## ACKNOWLEDGEMENTS

We thank the Department of Zoology, the Wellcome (WT096645MA and WT107060MA) for generous support, Malcolm Burrows for encouragement, and Gary Struhl and David Strutt for stocks and constructive criticisms.

## Supplementary Figure Legends

**Figure S1.**
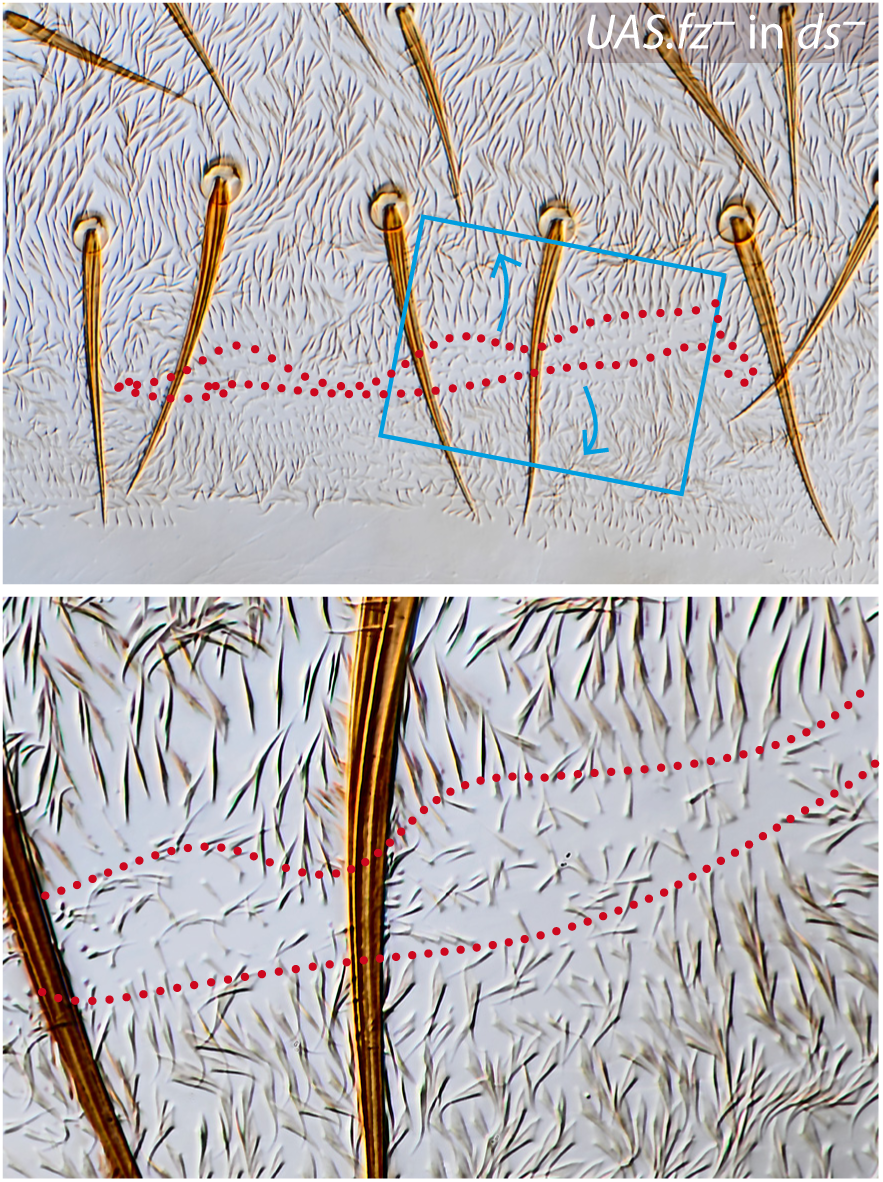
*fz*-overexpressing clone in the P compartment of a *ds^-^* fly. Hairs point outwards from the clone with range of 2–7 cells. Cells of the clone are marked with *pawn*, and outlined in red dots. Blue arrows indicate orientation of hairs. Blue box is enlarged below.

**Figure S2.**
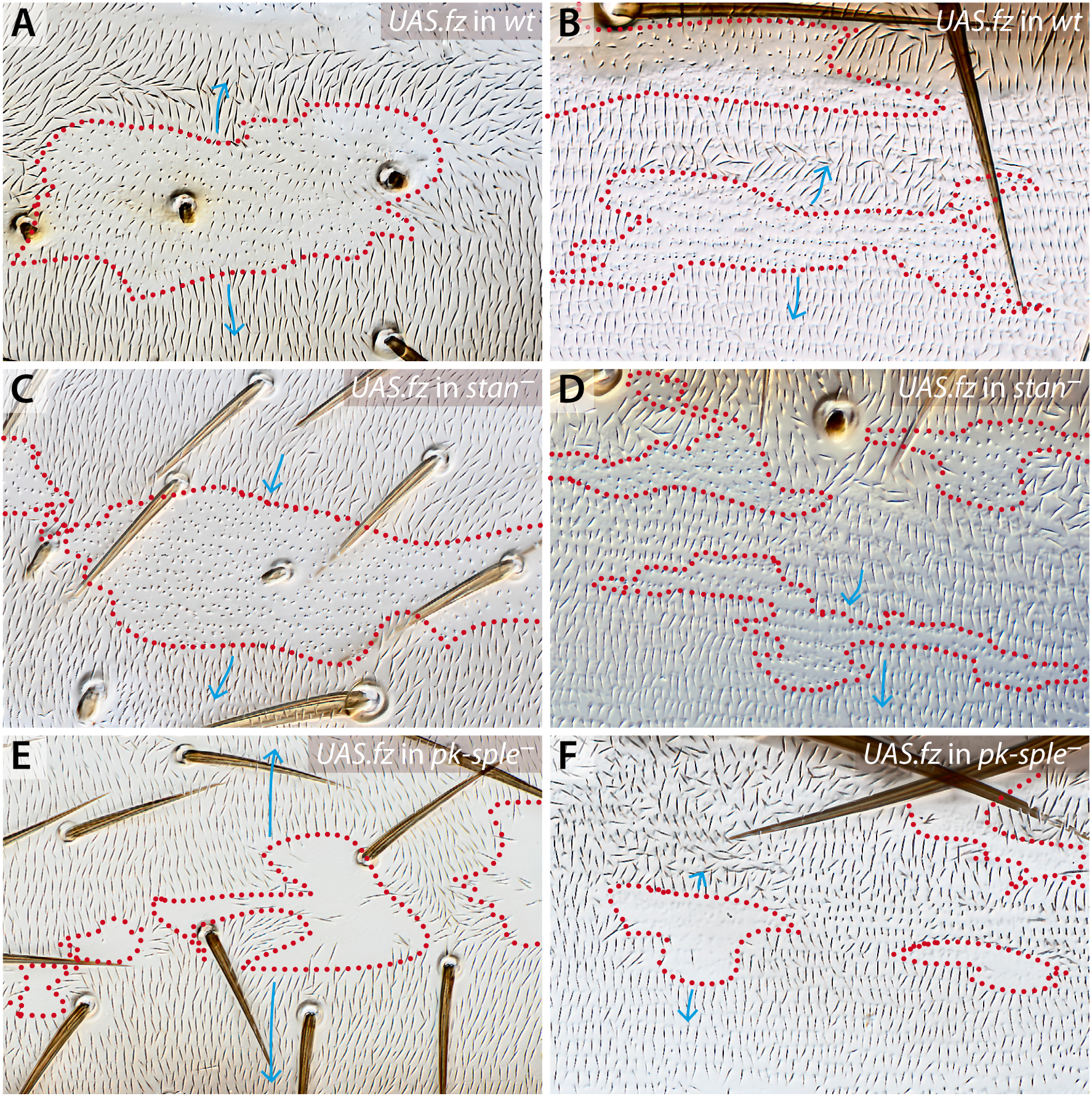
The effects of *fz*-overexpressing clones on various genetic backgrounds in the A and P compartments —compare with Figure 2. A compartments (**A**, **C** and E), P compartments (**B**, **D** and **F**). The clones polarise responding wildtype cells outwards in both compartments (**A** and **B**). This effect is blocked when the Stan system is broken (*stan^-^*) (**C** and **D**). In a *pk-sple^-^* background the sign is also outwards but the range of repolarisation is strongly reduced in the A compartment (**E** and **F**). Clones are variously marked, see Genotypes in Materials and Methods.

**Figure S3.**
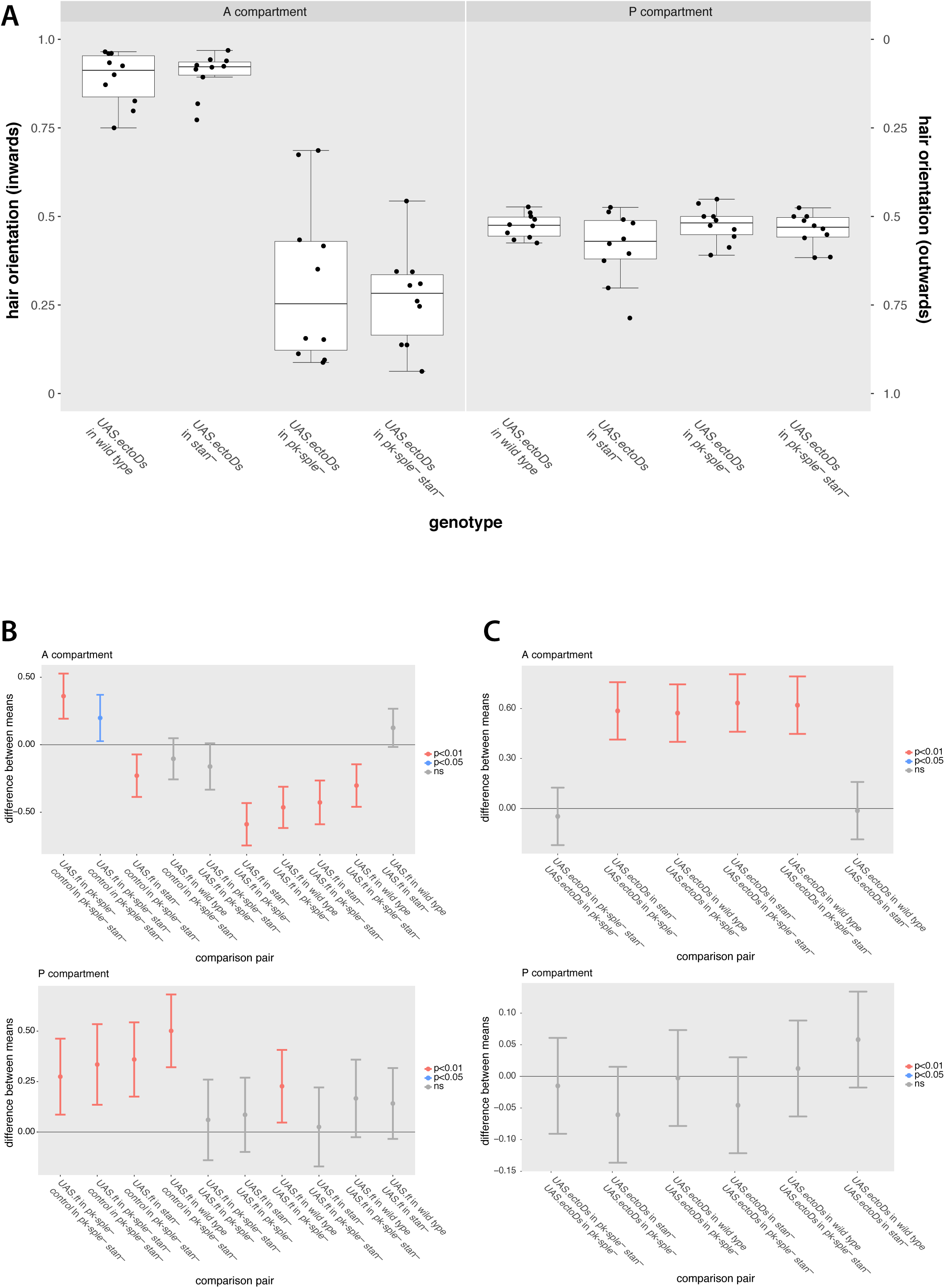
Results of similar experiments to those in Figure 3, but here the clones were overexpressing the ectodomain of Ds. The results are comparable with those of Figure 3 in the A compartments (although of the opposite sign to *ft*-overexpressing clones, as expected (Casal et al., 2006). None of the clones had significant effects in the P compartment — this lack of response is most simply explained by high ambient level of Ds in P, which is suggested by *ds.LacZ* expression (Casal et al., 2002). A response was visible in flies that lack *four-jointed* (*fj*) (data not shown), which increases the range of signalling by the Ds/Ft system (Casal et al., 2006). One-way Anova with post-hoc Tukey HSD analysis showing levels of significance for Figure 3 and S3, below (vertical lines are the 95% confidence intervals).

**Figure S4.**
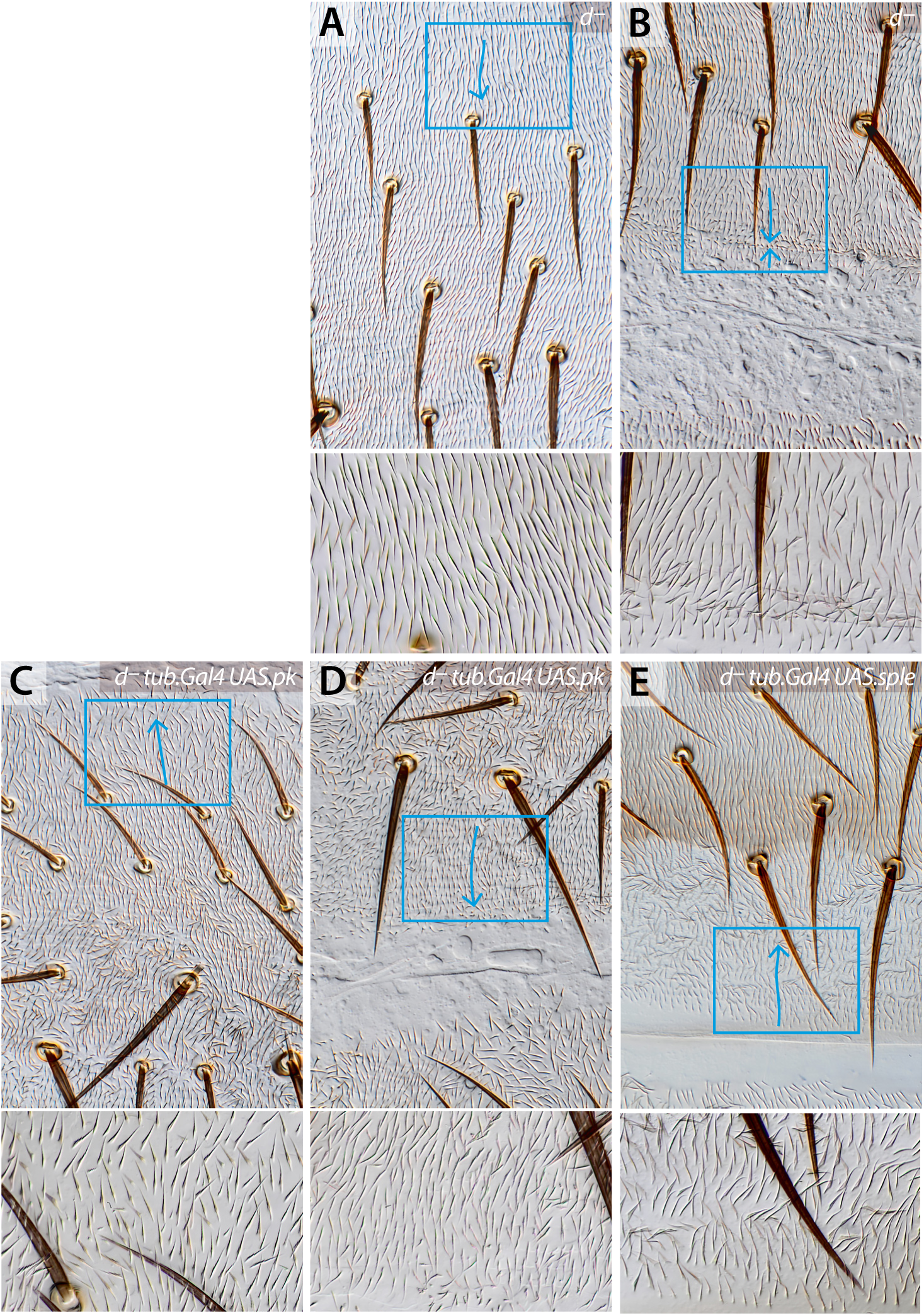
The effects of overexpression of *pk* and *sple* in *d^-^* flies. In this background the effects of extra Pk are as in *ft^-^ d^-^* flies: the anterior part of the A compartment points forward and the polarity of the P compartment is “rescued” (compare **C** and **D** with **A** and **B**; see Figure 4). However extra Sple increases the area of anteriorwards polarity in the P compartment (compare **E** with **B**; see Figure 5).

**Figure S5.**
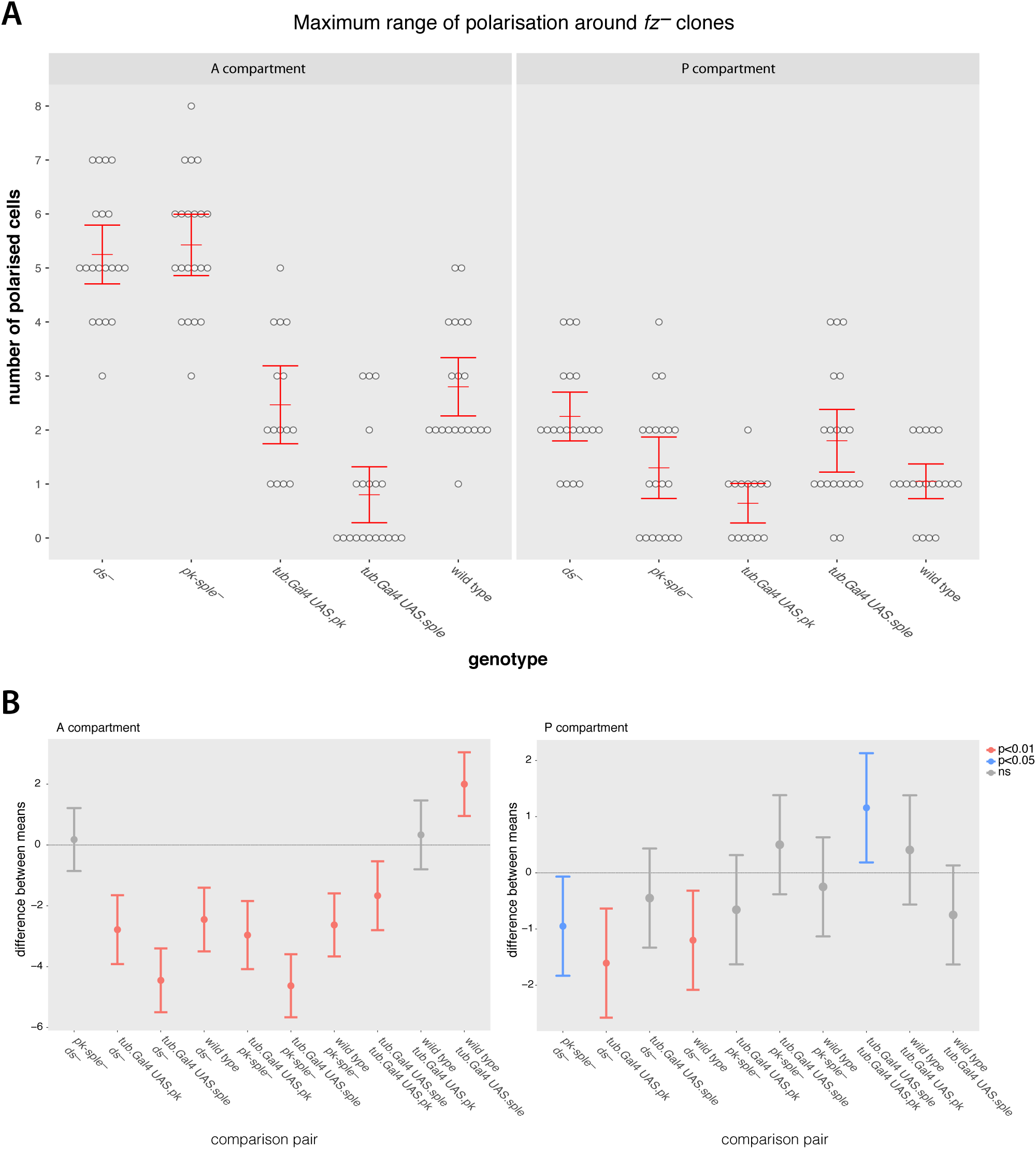
Range measurements for *fz*-expressing clones in wildtype and flies with a broken Ds/Ft system (*ds^-^*). For each clonal perimeter the maximum number of cell rows showing an induced polarity change was measured. Below are the results of one-way Anova with post-hoc Tukey HSD analysis.

**Figure S6.** Ventral cuticle of the abdominal segments stained for lacZ; **A**, *ds.lacZ* expression; **B**, *ds.lacZ* expression in *pk-sple^-^;* **C**, *fj.lacZ* expression; **D**, *fj.lacZ* expression in *pk-sple^-^*. Red dots delineate the approximate boundaries between the A and the P compartments. Arrows indicate the orientation of cell hairs in the pleura.

